# Modified *E. coli* strains enhance baculovirus production by elimination of aberrant transposition events

**DOI:** 10.1101/2021.01.27.427812

**Authors:** Carissa Grose, Chelsea Wright, Jennifer Mehalko, Dominic Esposito

## Abstract

Baculovirus technology has been the most commonly used expression system for insect cells both due to its potential to generate a large amount of recombinant protein as well as the benefit of post-translational modifications. The most commonly used system to generate recombinant baculoviruses is the Tn7 transposition-based technology known as Bac-to-Bac. Although improvements have been made to this system to further improve quality and reproducibility of baculovirus production, recent data suggests that improved strains still have potential issues with contamination of non-recombinant baculovirus caused by improper transposition into a Tn7 site in the *E. coli* chromosome. Here we describe a new option for alteration of the *E. coli* genome to completely block the native Tn7 attachment site, leading to far fewer false positive bacmid colonies being selected and eliminating all risk of non-recombinant baculovirus production.

## Introduction

The most common production method for recombinant baculovirus used for protein expression in insect cells is the Bac-to-Bac system and its derivatives (1–3). This system utilizes an *E. coli* host to generate *Autographa californica* multicapsid nucleopolyhedrovirus (AcMNPV) bacmids containing the gene of interest via Tn7-based transposition. To generate recombinant baculoviruses in such systems, an expression plasmid containing the gene of interest flanked by Tn7 right and left arms is transformed into the bacmid-containing strain (DH10Bac in the original system, or modified strains like DE26 for the newer B2F system), and the expression cassette between the Tn7 arms is transposed into a specific attTn7 attachment site on the bacmid by the transposition proteins TnsABCD located on the resident helper plasmid.

In addition to the attTn7 site on the bacmid, there is also an *E. coli* chromosomal attTn7 site located at the end of the essential *glmS* gene. It has been reported (4,5) that some percentage of the time, the expression cassette is shuttled to this genomic location rather than the desired bacmid transposition site leading to non-recombinant bacmid clones. The construction of the B2F strain, DE26, was an attempt to solve this problem by blocking the chromosomal attTn7 site with a mini-Tn7 insertion which included a chloramphenicol acetyltransferase (CAT) resistance marker and fully functional Tn7 left and right arms. Similar strains have been produced previously for the same purpose using analogous approaches. The transposition block was put into place via transposition and greatly reduced the number of false positives that were selected during the bacmid screening process (2). The presence of the mini-Tn7 at this location provided target immunity, disallowing additional Tn7 transposition into the neighboring regions (6). However, recent data from our laboratory shows that the functional Tn7 arms in the blocking construct still permit a low level of transposon mobility and we observe that the blocking cassette will occasionally move out of the chromosomal blocking location and back into the bacmid. This not only leaves the chromosomal location open to insertion of the recombinant gene, but also blocks the recombinant gene from bacmid insertion, creating a situation where some portion of the purified bacmid pool was carrying the CAT cassette rather than the intended gene of interest. The problem is further exacerbated by the standard process to verify insertion of the gene of interest—PCR analysis of the flanking regions around the bacmid insertion site produced false positive results since the blocking cassette, like the real recombinant region of interest, is flanked by Tn7L and Tn7R regions. Overall this issue leads to the possibility of mixed populations of bacmid, some of which may produce non-recombinant baculovirus that does not express the gene of interest. To address this issue, we devised two new blocking methodologies and constructed two new strains: DE77 where the TnsD binding region of the *glmS* gene sequence was mutated to eliminate transposition, and DE95 where only the Tn7 right arm was inserted at the chromosomal attTn7 target site (Fig 1). Previous literature had suggested that the Tn7R arm alone would be enough to provide target immunity (7).

**Figure 1:**
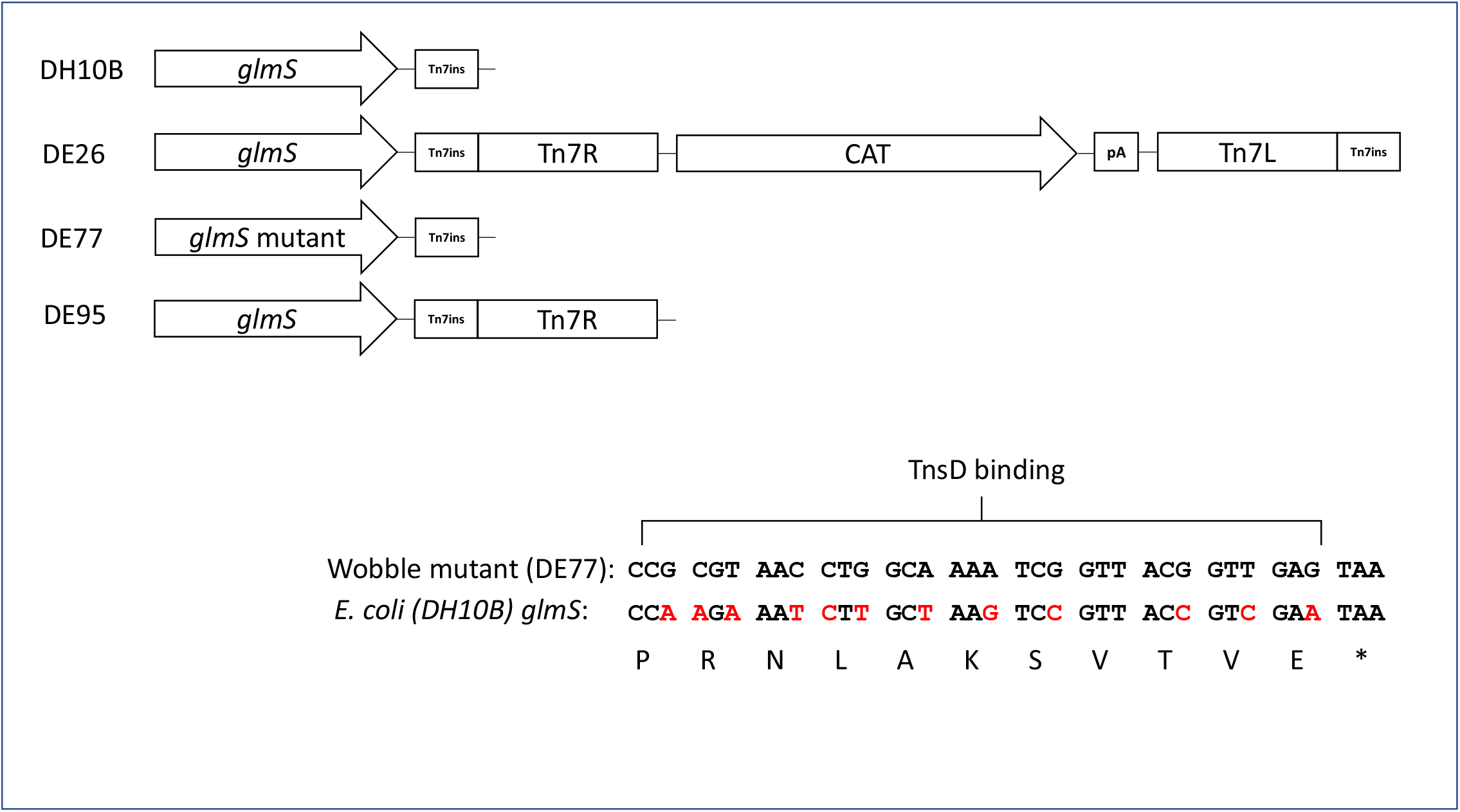
Schematic of *E. coli* strains used in this work. Shown are the regions around the Tn7 insertion site in the four strains discussed in this paper. Abbreviations are *glmS*, E. coli glutamine--fructose-6-phosphate aminotransferase; Tn7R, right Tn7 transposon end; Tn7L, left Tn7 transposon end; pA, SV40 polyadenylation sequence; CAT, chloramphenicol acetyl transferase; Tn7ins, insertion sequence for Tn7 downstream of the attTn7 site. The inset shows the sequence of the 3’ end of the E. coli *glmS* gene and the DNA changes made in the DE77 mutant.

### Recombineering Cassettes and Strain Construction

The recombineering cassettes used to generate DE75 and DE94 strains were constructed with two ~300bp arms homologous to the regions adjacent to the desired insertion location (scheme outlined in Fig. 2). For the mutant *glmS* of DE75, flanking oligos were used to alter the codon sequence of the TnsD binding region (final 11 amino acids), leaving the protein sequence intact (see Fig. 1, inset). For DE94, the Tn7R arm was amplified from the mini-Tn7 cassette of DE26. The homology arms, blocking sequences, and CAT-SacB dual selection marker from pELO4 (a generous gift from Don Court, NCI-Frederick) were assembled by isothermal assembly with 40bp of homology to the neighboring amplicon. Marker removal cassettes were generated by amplifying ~300bp fragments from either side of the CAT-SacB and assembling them by overlap PCR with 40bp of homology for a final ~600bp cassette. DE75 and DE94 were generated by λ RED recombination. The recombineering strain for this work was prepared by transforming *E.coli* DH10B with the λ RED plasmid, pKD46 (8). An ampicillin-resistant colony was then grown in LB with 100 ug/ml ampicillin overnight at 30°C and this seed culture was used to inoculate a fresh culture at 30°C with shaking to OD_600_=0.2. A final concentration of 0.1% arabinose was then added to induce the λ recombination genes. Culture growth continued to an OD_600_ of 0.5 and cells were made electrocompetent. Linear insertion cassettes (100 ng) were separately transformed into 50ul freshly prepared DH10B/pKD46 electrocompetent cells. Colonies were selected on LB with 30 ug/ml chloramphenicol and 100 ug/ml ampicillin followed by PCR confirmation of the insertion.

**Figure 2:**
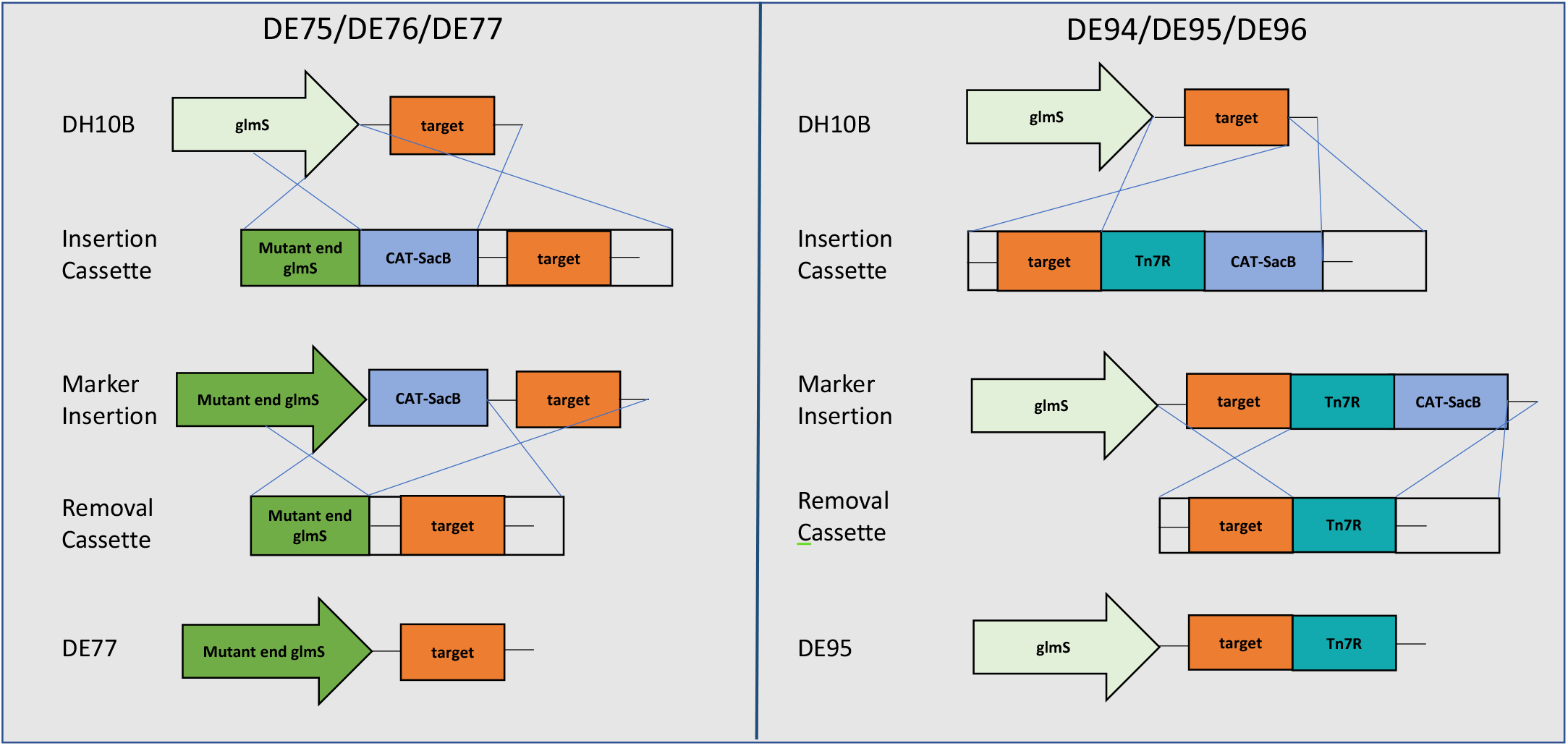
Recombineering strategy for construction of new blocking strains. The recombineering scheme for generating mutant strains and removing selection cassettes is shown. The CAT-SacB selection cassette contains a positive selection marker (CAT, chloramphenicol acetyltransferase) and a negative selection maker (SacB, levansucrase). Crossovers during recombineering are noted by blue lines.

A single positive culture for each strain was then made electrocompetent after arabinose induction as described above. The CAT-SacB selectable marker was excised using 100ng of the 600bp removal cassettes and colonies were selected on LB no salt/6% sucrose/100 ug/ml ampicillin. Final strains (Table 1) were validated by PCR amplification of the altered region and confirmed by Sanger sequencing. After confirmation of the genome modifications, the pKD46 plasmid which carries the repA101ts temperature sensitive origin was removed by two overnight growths on LB plates at 37°C. Diluted colonies were then streaked on LB/ampicillin plates to confirm the ampicillin sensitivity via loss of pKD46.

**Table 1:** Strains used/generated in this work. Strains labeled as “+ Helper” contain the R982-X01 helper plasmid while those with “+ Helper,+Bacmid” contain the R982-X01 helper and the standard B2F bacmid construct. Detailed descriptions of the construction of DE25 and DE26 can be found in (2).

|  | **Base Strain** | **+ Helper** | **+Helper +Bacmid** |
| --- | --- | --- | --- |
| CAT block | DH10B | DE25 | DE26 |
| *glmS* mutant | DE75 | DE76 | DE77 |
| Tn7R block | DE94 | DE96 | DE95 |

The B2F helper plasmid, R982-X01 (2) was transformed into the new strains and colonies were selected on LB with 12.5 ug/ml tetracycline to create the DE76 and DE96 strains. Finally, a bacmid lacking the chitinase and cathepsin genes (9,10) was transformed into these new strains and colonies were selected on LB with 12.5 ug/ml tetracycline and 50 ug/ml kanamycin to generate the final DE77 and DE95 strains.

### Stability of Blocking Regions, Blocking Efficiency, and Recombinant Gene Insertion Efficiency

The mobility of the blocking regions of each strain was checked by growing the strains overnight, preparing alkaline lysis DNA (which will contain both bacmid and genomic DNA), and PCR analysis of the attTn7 regions of both the bacmid insertion site region and the *E. coli* chromosomal attTn7 region. In Fig. 3, DE77 and DE95 blocking sequences are stable as there is no evidence of unexpected bands by PCR. However, the DE26 bacmid region showed unexpected amplicons. The size of the 1.9 kB PCR fragment observed was consistent with transposition of the blocking cassette from the chromosome back into the bacmid attTn7 site—this was confirmed by Sanger sequence analysis of the PCR fragments which showed a clean transposition of the blocking cassette into the bacmid.

**Figure 3:**
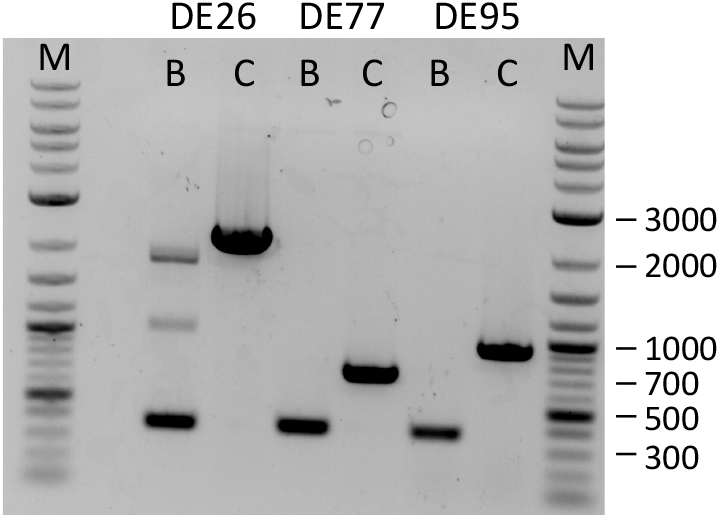
PCR confirmation of stability of blocking sequences. For each strain, PCR amplicons of the bacmid attTn7 region (B) and the *E. coli* chromosomal attTn7 region (C) are shown. Lanes labelled M are DNA markers (sizes noted in bp). Empty bacmid regions are expected to produce a 375bp fragment while stable chromosomal blocking regions should yield 2.2 kbp (DE26), 778 bp (DE77), and 945 bp (DE95) bands. Movement of the CAT mini-Tn7 in DE26 to the bacmid would yield an amplicon of 1.9 kbp.

To test the efficiency of the three *E.coli* sequences at blocking insertion of a recombinant gene, we transformed a B2F (non-replicative) expression plasmid, R951-M01-382 (2), into strains (DE25, DE76, DE96) containing the helper plasmid, but lacking bacmid DNA. We plated 5% and 25% of the transformation mixtures (more than 10x and 50x the number of cells in a usual bacmid production) in order to detect any small amount of growth, as a strain with ideal genomic blocking should yield very few or no colonies in the absence of the bacmid. As Table 2 shows, DE25 and DE96 plates had very few colonies while DE76 plates had thousands, indicating that the mutation of *glmS* was insufficient to prevent binding of the TnsD protein. Several colonies from each plate were analyzed by PCR to see if the recombinant gene could be detected on the chromosome. None of the rare colonies for DE25 or DE96 showed such insertions, while 100% of colonies from the DE76 plate were positive for insertions, further confirming the ineffective blocking of that region by the mutations.

**Table 2:** Colony counts from ampicillin-containing plates after transformation and plating with the indicated amount of a 0.5 ml transformation reaction.

|  | 5% | 25% |
| --- | --- | --- |
| DE25 | 3 | 8 |
| DE76 | ~2000 | ~4000 |
| DE96 | 0 | 1 |

The same B2F expression plasmid was transformed into the three bacmid strains to see how efficiently the gene of interest would be transposed into the bacmids and whether the blocking region would move out of its intended location in the presence of the bacmid. Six colonies were selected from each bacmid plate for overnight liquid culture growth. DNA was prepared by alkaline lysis and each was used as a template for PCR analysis of both the bacmid and chromosomal attTn7 regions. In the selected samples, 100% of the DE26 and DE95 bacmids had the recombinant gene present in the bacmid and still maintained the expected genomic block of the E. coli attTn7 site. In the DE77 samples, 1 of the 6 bacmids lacked insertion into the bacmid and 3 of the 6 had insertion of the recombinant gene in the chromosome, again highlighting the failure of the DE77 blocking methodology.

## Conclusions

While the DE26 B2F bacmid strain was mostly successful at blocking transposition into the chromosomal attTn7 site, it was still capable of transposing back out of this location and into the bacmid attTn7 site. This poses a significant risk for high-throughput applications where a mixture of recombinant and non-recombinant baculovirus can easily be created. The mutated TnsD binding site of DE77 was insufficient to block chromosomal insertion, but the Tn7R cassette in DE95 both blocked the undesired insertion as well as allowed high efficiency insertion into the bacmid. We anticipate that this new strain will improve expression levels for some recombinant proteins as the purified bacmid DNA will more reliably contain the gene of interest, and will also eliminate the possibility of selection for non-recombinant baculovirus which could affect long-term stability of baculovirus cultures.

## Acknowledgments

This project has been funded in whole or in part with Federal funds from the National Cancer Institute, National Institutes of Health, under contract number HHSN261200800001E. CW was supported by a National Cancer Institute Summer Internship in Biomedical Research (SIP) fellowship. The content of this publication does not necessarily reflect the views or policies of the Department of Health and Human Services, nor does mention of trade names, commercial products, or organizations imply endorsement by the U.S. Government.

## Author Contributions

CG: Conceptualization, Methodology, Validation, Investigation, Resources, Writing-Original Draft; CW: Investigation; JM: Conceptualization, Methodology; DE: Conceptualization, Writing-Review & Editing, Supervision, Project Administration, Funding Acquisition.

